# Cross-species imputation and comparison of single-cell transcriptomic profiles

**DOI:** 10.1101/2023.10.19.563173

**Authors:** Ran Zhang, Mu Yang, Jacob Schreiber, Diana R. O’Day, James M. A. Turner, Jay Shendure, Christine M. Disteche, Xinxian Deng, William Stafford Noble

## Abstract

Cross-species comparison and prediction of gene expression profiles are important to understand regulatory changes during evolution and to transfer knowledge learned from model organisms to humans. Single-cell RNA-seq (scRNA-seq) profiles enable us to capture gene expression profiles with respect to variations among individual cells; however, cross-species comparison of scRNA-seq profiles is challenging because of data sparsity, batch effects, and the lack of one-to-one cell matching across species. Moreover, single-cell profiles are challenging to obtain in certain biological contexts, limiting the scope of hypothesis generation. Here we developed Icebear, a neural network framework that decomposes single-cell measurements into factors representing cell identity, species, and batch factors. Icebear enables accurate prediction of single-cell gene expression profiles across species, thereby providing high-resolution cell type and disease profiles in under-characterized contexts. Icebear also facilitates direct cross-species comparison of single-cell expression profiles for conserved genes that are located on the X chromosome in eutherian mammals but on autosomes in chicken. This comparison, for the first time, revealed evolutionary and diverse adaptations of X-chromosome upregulation in mammals.

## 1 Introduction

The magnitude of a gene’s expression may vary across species, and this variation may contribute to or be representative of morphological or trait evolution [1, 2]. Thus, comparing gene expression profiles across species has the potential to offer valuable insights into a wide range of questions related to, for example, which genes have adapted to new regulatory machineries and functions during evolution, and how gene expression changes when a gene moves to a different chromosomal context or when a gene’s copy numbers varies across organisms (e.g., genes located on two sets of autosomes in chicken and on the single X chromosome in male mouse). Furthermore, understanding transcriptional differences between model organisms and humans will greatly enhance our ability to transfer insights gained from model organism studies into a human context.

Previous studies have compared transcriptional differences among organisms based on bulk and single-cell gene expression measurements. However, comparison of bulk gene expression profiles across species [3–6] may not fully capture cellular heterogeneity and may suffer from imbalanced cell type composition of tissues across species [7]. Recently, single-cell profiles have been widely generated to capture cell-specific expression profiles and mitigate the issue of uncaptured cell heterogeneity in bulk samples, but direct transcriptional comparisons are difficult because of the challenge of matching cells across species. Moreover, single cell measurements suffer from sparsity, noise, variable sequencing depth, and potential batch effects. As a result of these limitations, current studies instead perform cross-species matching or comparison at the cell type level [8–12]. Unfortunately, this approach requires accurate cell type calling and matching across species and, similar to bulk comparisons, fails to take into consideration single-cell variability.

More importantly, although single cell data are widely available for certain model organisms, such as mouse, single-cell expression atlases in human and non-model-organisms are still far from complete, due to limitations of sample availability and accessibility (e.g., it is difficult to retrieve human brain samples, as well as fetal or pediatric tissues, especially in disease conditions). Although multiple methods have been developed that focus on aligning single cells across species [8, 10, 13, 14], these methods are not able to make cross-species comparison at single-cell resolution, nor can they make predictions about evolutionary changes when certain experimental measurements (e.g., a particular cell or tissue type of a species) are not available. Because of these limitations, studies of transcriptional evolutionary patterns are restricted to tissues or cell types in species that have existing, high-quality measurements.

To address these challenges, new computational methods are needed that can (1) predict gene expression profiles for missing cell types and biological contexts and (2) directly compare expression profiles across species at single-cell resolution, without relying on external cell type annotations. Although several deep learning-based methods enable cross-species prediction of perturbation effects from high-throughput screens, these methods rely on discrete cell type labels and are not designed to predict and compare cellular gene expression in wildtype physiological conditions across species [7, 15, 16]. We hypothesize that a neural network model that can decompose single cell profiles into species factors and cell factors invariant of species will allow us to make single-cell, cross-species prediction and comparison by swapping the species factor corresponding to each cell.

We are motivated in part by the study of sex chromosome evolution, which would benefit from methods for single-cell comparison and prediction of gene expression across species. In mammals, males have a single X chromosome and a gene-poor Y chromosome, whereas females have two X chromosomes. X chromosome upregulation (XCU) has been proposed to evolve in response to gene loss due to Y-chromosome degeneration during sex chromosome evolution [17–22]. XCU increases expression of many X-linked genes to balance gene expression between the single X chromosome and two sets of autosomes in diploid male XY cells. In female XX cells, X inactivation silences one of the two X chromosomes in females to avoid hyperactivation. Evolutionary studies have shown that the X chromosome of eutherian mammals has arisen from ancestral autosomes in two major steps: first, an autosome gave rise to the so-called X conserved region (XCR) in the ancestor of both eutherian and metatherian mammals [20]. Today, the XCR is represented by the whole X chromosome of marsupials (e.g., opossum) and corresponds to about two thirds of the eutherian X chromosome. The latter acquired the so-called X added region (XAR) by translocation of autosomal pieces to the XCR, which resulted in the larger conserved X chromosome of eutherian mammals (e.g. human, mouse, rat). Chromosomes that demonstrate homology to the XCR and XAR can be identified in marsupials (X chromosome and parts of autosomes 4 and 7) and in birds (parts of autosomes 1 and 4 in chicken) (Figure 1D). While XCU has been clearly demonstrated in Drosophila and C. elegans, the mechanisms of XCU in mammals are still debated [23–27]. Importantly, which X-linked genes are upregulated and at which levels at each evolutionary step is still unclear. This is largely because most studies have relied on transcriptomic approaches to compare expression between groups of X-linked and autosomal genes. However, direct expression comparisons of evolutionarily conserved genes before and after becoming X-linked are limited or impeded by different data normalization methods, and by the lack of appropriate samples to directly compare species across tissues and cell types [4, 5].

**Figure 1:**
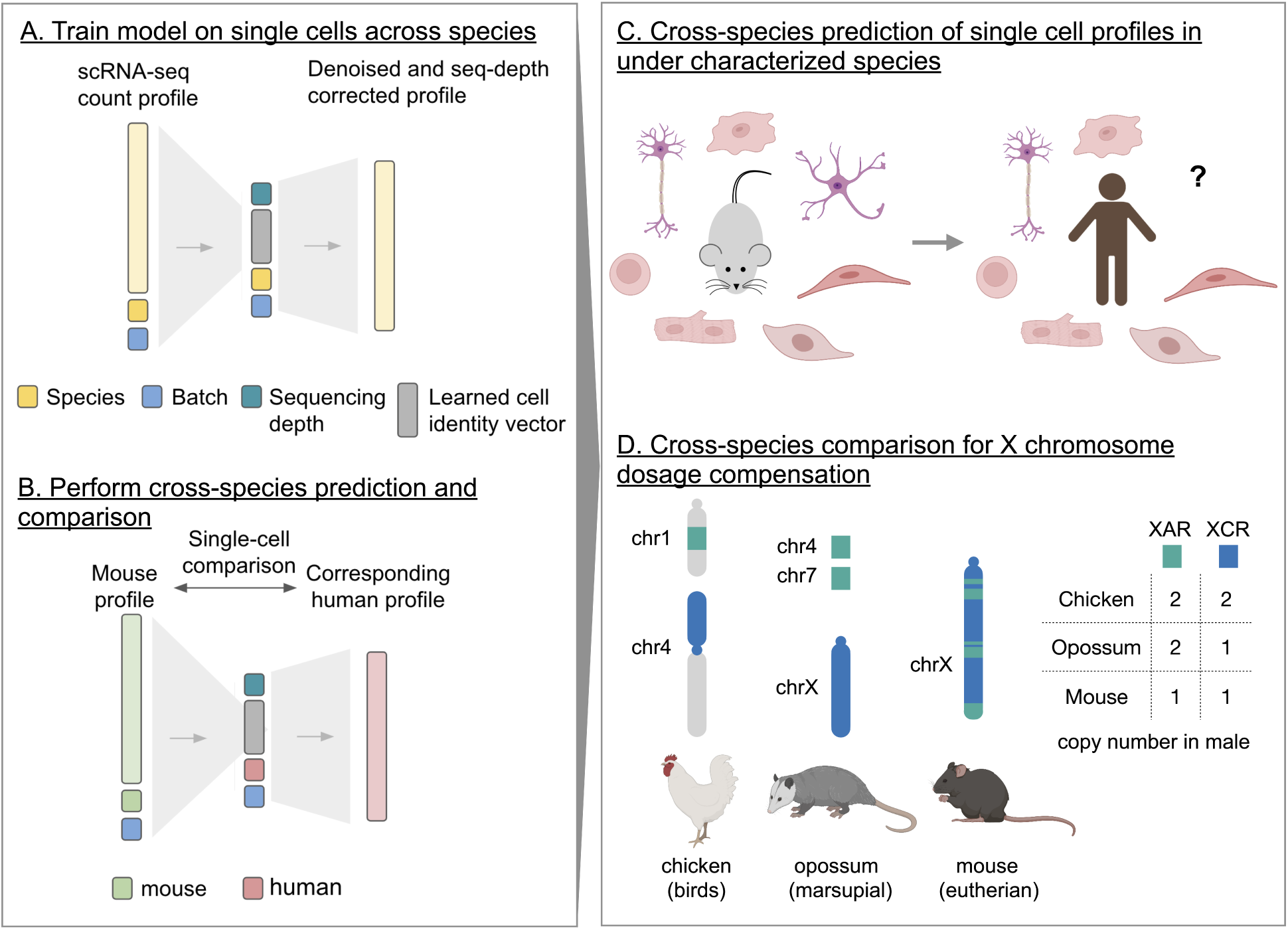
Icebear’s cross-species prediction and comparison framework. (A) Icebear is trained on single-cell RNA-seq profiles across species. Specifically, Icebear uses conditional variational autoencoders that predict cellular profiles from a combination of species and batch factors, as well as species-invariant cell factors. (B) Once the model is trained, we can swap the species factor to predict a (e.g., mouse) cell’s corresponding profile in another species (e.g., human). (C) Icebear can be used to predict single-cell profiles in species (e.g. human) where the corresponding cell type has not been profiled. (D) Icebear can also perform cross-species comparison at the single-cell level, revealing X chromosome up-regulation patterns during evolution.

Here we propose a deep learning model, Icebear, that induces a non-sparse version of single-cell expression data and performs cross-species prediction and comparison at single-cell resolution. Using several publicly available datasets, we demonstrate that Icebear is able to integrate single-cell expression profiles across species, batch and tissue types, and predict single-cell profiles in missing cell types across species. In addition, we show that, on the basis of mouse data, Icebear can accurately predict transcriptomic alterations in human Alzheimer’s disease (AD) versus control samples, thereby enabling the transfer of knowledge from singlecell profiles in mouse disease models to human. After several cross-species validation experiments based on public datasets, we applied Icebear to predict and compare gene expression changes across eutherian mammals (mouse), metatherian mammals (opossum) and birds (chicken), using our in-house generated sciRNA-seq profiles with minimal cross-species batch effects. By doing so, Icebear reveals gene expression pattern shifts across species that support the existence of mammalian XCU and suggest the extent and molecular mechanisms of XCU vary among mammalian species and among X-linked genes with distinct evolutionary origins.

## 2 Methods

### 2.1 Multi-species single cell profile generation

Mixed-species scRNA-seq data were generated by a three-level single-cell combinatorial indexing approach (sci-RNA-seq3) [28]. Adult brain and heart from both male mouse and chicken were purchased from BioChemed Services, and male opossum adult brain was provided by J. Turner (MRC, UK). The data were collected by indexing cells from each species by reverse transcriptase barcoding and then processing them jointly, in which case the species identity of each cell was known based on the sequence barcode.

### 2.2 Assigning species labels to cells and mapping reads

Mapping reads from sci-RNA-seq3 experiments to profile hundreds of thousands of single cells from multiple species’ samples is complicated by the possibility of cells from different species entering the same doublet during sample preparation and three-round combinatorial cell barcoding. Accordingly, our pipeline begins by mapping each read to multiple species and retaining only reads that map uniquely to a single species. This step allows us to detect and remove species-doublet cells with reads from more than one species.

The detailed protocol is as follows:

1. For a given sample, create a multi-species reference genome by concatenating the reference genomes of all the species used in that sample.
2. Map all of the reads to the multi-species reference, retaining only reads that map uniquely. We used the STAR aligner [29] with the following parameters: --outSAMtype BAM Unsorted –outSAMmultNmax 1 --outSAMstrandField intronMotif --outFilterMultimapNmax 1.
3. Remove PCR duplicates.
4. Eliminate any read that maps to an unassembled scaffold, mitochondrial DNA, or any locus that is marked as a repeat element by RepeatMasker (http://www.repeatmasker.org¿). The repeat elements by RepeatMasker were retrieved from UCSC genome browser [30], with the exception of opossum, where we ran RepeatMasker to generate them. Repeat elements were removed using BEDtools [31].
5. For each cell, count the total number of remaining reads that map to each of the three species.
6. If the sum of the second- and third-largest counts is greater than 20% of all counts, then mark the cell as a species-doublet and eliminate it.
7. Label the remaining cells according to their generating species.

Having identified the species origin for each cell, we then re-map the reads associated with each single-species cell only to its corresponding species, retaining only the reads that map uniquely within that species.

The parameters used for this step were --outSAMtype BAM Unsorted --outSAMmultNmax 1 --outSAMstrandField intronMotif.

The re-mapping is run using the pipeline developed by the Brotman Baty Institute (BBI) (https://github.com/bbi-lab/bbi-sci/). The first two steps of mapping multi-species reference were modified from their pipeline.

The reference genomes and annotations were from the BBI, using assembly GRCm38 for Mus musculus, ASM229v1 for Monodelphis domestica, and GRCg6a for Gallus gallus (Ensembl release 99) [32]. The genes were also filtered to match the ones used by BBI.

### 2.3 Reconciling orthology relationships

To simplify the model and focus on the most straight-forward cross-species transcriptional changes, we would like to establish one-to-one orthology relationships among genes in the various species included in our study. The Biomart resource [33, 34] at Ensembl reports orthology relationships between genes in a many-to-many fashion, each with an associated percent identity score. To reduce this data resource to a one-to-one mapping, our approach takes into account the following considerations.

- We use the percent identity score to resolve ambiguities. If, for example, two mouse genes are mapped to two opossum genes with four edges, then we select the two edges that create a one-to-one mapping with maximal score.
- In some cases, transitive relationships can be used to fill in missing edges. For example, if gene *A* in human has an ortholog *B* in mouse, *B* has an ortholog *C* in chicken, but *A* has no ortholog in chicken, then we can add an edge from *A* to *C*.

Formally, we represent our problem using an undirected graph *G* in which vertices *V* are genes and edges *E* represent orthology relationships. Each vertex *v ∈ V* has an associated species label *s*(*v*) *∈ {*human, mouse, rat, opossum, chicken*}*. Our goal is to eliminate edges from this graph so as to ensure that every node is connected to at most one node in each of the other species This is done by building a graph *G^t^* = *{V ^t^, E^t^}* such that *V ^t^ ⊂ V* , as follows. First, we reconcile mouse with each other species by creating a one-to-one mapping between genes in the two species. We do this in a greedy fashion for two species *A* and *B* by ranking all edges connecting *A* and *B* (i.e., all “high confidence” edges in Biomart) in decreasing order by percent identity and then adding an edge to *G^t^* if and only if neither of its corresponding vertices already has an associated accepted edge. Second, we fill in transitive relationships. This is done by creating a one-to-one mapping between genes, using the same greedy algorithm as before, but for each pair of non-mouse species. We then search for “transitivity” triangles (*A, B, C*) that fulfill the following criteria:

- *s*(*A*) = human
- *s*(*B*) *I*= human and *s*(*C*) *I*= human
- *s*(*B*) *I*= *s*(*C*)
- The edge from *A* to *B* is in *G^t^*.
- *G^t^* does not contain any edge connected to *C*.
- The one-to-one mapping between *s*(*B*) and *s*(*C*) connects *B* to *C*.

In this case, we add to *G^t^* the edge connecting *A* to *C*. This transitivity step is done iteratively over the non-mouse species in order of evolutionary distance; i.e., the species *B* and *C* are considered in the following order: (mouse, opossum), (mouse, chicken), (opossum, chicken). For each of the resulting triples of species, transitivity triangles are selected in a greedy fashion by sorting the triangles in decreasing order by the percent identity associated with the edge connecting *B* to *C*.

At the end of this process, in the graph *G^t^* each mouse vertex has at most one connected neighbor in each of the non-mouse species. Note that we make no attempt to ensure that the pattern of orthology relationships respects the species tree. Thus, in principle a mouse gene might have an ortholog in chicken but none in opossum. This is possible in the case of a gene deletion event along the opossum lineage. This process yielded 10,030 genes with orthologs shared across the three species, 70 of which were added through transitive relationships.

This process yielded 10,030 genes with orthologs shared across the three species, 70 of which were added through transitive relationships.

### 2.4 Data preprocessing

#### 2.4.1 In-house multi-species dataset

To perform cross-species prediction and comparison, we performed the following preprocessing steps:

1. Retain genes that have orthologs shared across all species.
2. Remove all mitochondrial reads.
3. Remove cells with *<*200 UMIs.
4. Remove genes expressed in fewer than 50 cells across all the datasets.

This process resulted in a gene (n=9878) by cell (m=561340) matrix (Table 1), the median UMI is 450.

**Table 1:** Cell counts in the final dataset.

| species | tissue | batch1 | batch2 | batch3 |
| --- | --- | --- | --- | --- |
| mouse | brain | 33569 | — | — |
| opossum | brain | 2664 | 235178 | — |
| chicken | brain | 8927 | 167873 | — |
| mouse | heart | — | — | 17403 |
| chicken | heart | — | 28003 | 67723 |

#### 2.4.2 Public datasets

Gene expression profiles were downloaded from a multi-species primary motor cortex (M1) dataset [35], human and mouse cell atlases [36, 37] and an Alzheimer’s disease study [38].

For the M1 dataset, we used the expert-curated cell type annotations in each dataset, which contains 10 major cell type annotations that are further subdivided into 45 high-resolution cell type annotations. We only focused on cell types that are annotated as homologous between human and mouse based on expert curation. After applying the above gene filtering, mapping and cell filtering steps, 233,296 cells and 13,924 genes are retained for the downstream analysis.

For the human and mouse atlases, we trained Icebear on cells collected in the adult stage, which contains

288,886 cells and 12,367 genes. For the purpose of evaluation, cell types were determined based on expert annotations drawn from the original paper, and we only made predictions on cell types with *≥* 25 cells in both human and mouse. To make sure our predictions are not confounded by batch or donor effects, we only evaluated on the subset of cell types that exist in tissues that do not exhibit large donor effects in humans. To do that, we calculated Euclidean distances based on normalized pseudobulk profiles across all human adult tissue samples, and we retrieved tissues in which samples from different donors are closest to each other.

For the Alzheimer’s disease dataset, we retrieved single cells from human and 7-month-old mice. For the purpose of evaluation, we mapped cell type annotations from mouse to human based on the cell clusters called in the original paper, and we only validated on cell types with one-to-one mappings across species. No additional filtering was performed. This dataset was merged with the M1 dataset (with the same ortholog mapping approach above) to increase the cell numbers for model training. 440,689 cells and 12,474 genes are retained in the joint dataset.

Because the model takes raw counts as input, no further data normalization is needed.

### 2.5 Icebear cross-species prediction model

Our model reconstructs each scRNA-seq profile using a set of factors representing cell identity, tissue, species, and batch. The batch factor represents any knowledge of potential technical confounders, including studies, sequencing machines, or data generation batches. In the absence of prior knowledge of experimental batches, the batch factor is the same across all cells. Similarly, the tissue factor is kept constant if all cells are extracted from the same tissue type. These factors are concatenated and fed into a neural network to reconstruct the original gene expression profile. The model adapts a conditional variational autoencoder framework [10]. Species, tissue, and batch factors are encoded using one-hot encoding, while the model learns cell factors as *n*-dimensional vectors, where *n* is a hyperparameter to be tuned.

Intuitively, each gene expression count (*X*) in a specific cell is estimated based on three learned variables: the sequencing depth-corrected mean (*µ^t^*), dispersion of the negative binomial distribution (*r*), and logit of the dropout event (*p*). The reconstruction loss consists of the log likelihood of the raw count *x* and sequencing depth-corrected estimation using a zero-inflated negative binomial (ZINB) loss:

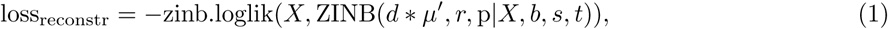

where *d* is the sequencing depth. The model also regularizes the cell embeddings by applying a KL divergence loss between the posterior distribution (*Q*) of cell embeddings (*z*) given batch (*b*), species (*s*) and tissue (*t*) factors and the prior distribution (*P* , standard multivariate normal distribution):

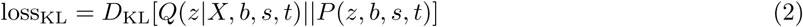

Thus, in each epoch we minimize the the following loss:

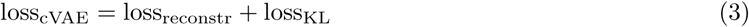

To further guide the alignment of cells across species, optionally, we adopted the idea of generative adversarial networks (GANs) [39], where we used a discriminator to distinguish cells from different species, and then we trained the model to fool the discriminator and learn cell embeddings invariant to species. Specifically, we divided the model training into two major iterative steps. In the first step, we trained a single-layer discriminator to predict species labels (*s*) based on cell embeddings, to distinguish species of origin from learned cell embeddings. In this step, the model aims to minimize the discriminator loss calculating the discrepancy between true species label (*s*) and predicted label (*D*(*z*)):

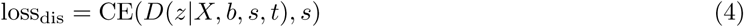

In the second step, we fixed the parameters in the discriminator and tried to optimize for reconstruction and fooling the discriminator by minimizing loss_cVAE_ *−* loss_dis_. The above steps are iterated per training epoch. This GAN option is provided as an option in the hyperparameter search.

In each prediction task, we held out all cells in a cell type or tissue in the target species as the test set, to mimic the actual use case where we would like to predict cellular profiles in an unseen context. This set of cells was not seen by the model during training and was used to evaluate prediction performance. For the rest of the cells, because our model does not rely on cell type annotations, we randomly assigned 10% to the validation set (with a cap of 20,000 cells), and the rest of cells were used as the training set. Each model was trained until the validation losscVAE stopped decreasing for 45 consecutive epochs. We then selected the model with minimum validation loss.

We selected model hyperparameters using a grid search strategy. We considered two hyperparameters: a Boolean indicating whether to include the discriminator layer or not, and the dimension of the bottleneck layer of 25, 50, 100. The number of hidden layers in the encoder and decoder are fixed to 2. Among these possible hyperparameter settings, we selected the model that yielded the largest LISI score [13] across cell embeddings learned from different species on the validation set.

To perform cross-species prediction on single cells, we first learned cell embeddings from cellular profiles in the known species and then concatenated the cell embeddings with the species factors of the target species to make cross-species predictions through the trained decoder architecture. This process produced sequencing depth-normalized and denoised single cell gene expression in the target species for each corresponding cell in the source species.

### 2.6 Cross-species prediction evaluation metric

To evaluate the prediction performance, we made use of expert-curated cell type labels and cell type matching across species, and we asked whether the model can accurately predict pseudobulk gene expression profiles of the missing cell types across species.

For each cell type, we held out its profile in the target species and made predictions using the corresponding cell type’s profile in the source species. Pearson correlation between the true and predicted pseudobulk profiles was used to assess the prediction performance (“prediction”). Pseudobulk profiles were calculated as the average of sequencing-depth-normalized gene expression profiles across cells, to put the true and predicted profile on the same scale.

We compared the prediction performance with three baselines.

1. Donor baseline (“donor baseline”): a “cheating” baseline that calculates the mean similarity across donors within the held-out test set. Here, we calculate the Pearson correlation of sequencing-depth normalized and log-transformed pseudobulk cell-type specific profiles between each pair of donors and report the mean correlation across all donor pairs.
2. Species baseline (“species baseline”): similarity between the original held-out cell type’s profile in human and the corresponding cell type’s profile in mouse. Genes are transferred to the target species through ortholog mapping and Pearson correlation between human and mouse sequencing-depth normalized and log-transformed pseudobulk cell-type specific profiles are reported.
3. Celltype baseline (“celltype baseline”): a “lazy” but close baseline that calculates the similarity between profiles of the held-out cell type and a similar cell type that has measurements in the target species. We first identify the cell type that has the most similar pseudobulk gene expression profile to the held-out cell type in the source species, calculated based on Euclidean distance on the sequencing-depth-normalized and log-transformed pseudobulk profiles. This baseline is created to ensure that our prediction captures more meaningful signals of the held-out cell type compared to guesses using a similar existing one.

To make a fair comparison with the performance of the donor baseline, we also calculated the mean of similarity measurements between our predictions with each donor in the test set (“indiv predictions”).

To measure the prediction error with regard to the original gene expression magnitude and variation, for each held out cell type, we calculated the relative prediction error based on pseudobulk gene expression 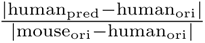.

### 2.7 Evaluation of Alzheimer’s disease profile prediction

To evaluate how well we can predict human AD profiles based on mouse models, we tried to place ourselves in the actual use cases when studying human diseases, which is to find differentially expressed genes and the direction and magnitude of expression alterations between AD and wild type (WT) in human samples. Because the predicted expression values in Icebear are denoised continuous values, it is not straightforward to statistically compare Icebear’s predictions with the sparse, count-based measurements in the original single-cell profiles. Therefore, we evaluated Icebear’s performance based solely on the direction and magnitude of gene expression changes in predicted AD vs WT profiles in human. These patterns are compared against the observed gene expression alternation patterns between the original AD versus WT profiles in human, based on Pearson correlation. Because studies of mouse disease models usually directly use the gene expression alterations in mouse as a proxy to understand human disease (when human samples are unavailable), we included gene alteration patterns in AD versus WT mouse as a baseline prediction, and calculated its correlation with gene expression alterations in human.

To capture both the direction and magnitude of gene expression alterations in AD and WT, we used the log2 fold change (log2FC) based on pseudobulk profiles (normalized to library size of 10,000) as the gene expression alteration measurement. To further avoid large, noisy log2FCs from genes with low expression values, we added 1 to all pseudobulk gene expression profiles before calculating log2FC.

### 2.8 Cross-species X-linked gene expression changes during evolution

To assess the pattern of XCU during mammalian sex-chromosome evolution, we first trained the model on all datasets and then compared gene expression changes between each pair of species by directly swapping the species factors. To do that, we used the trained Icebear model to retrieve cell embeddings based on single-cell profiles in mouse, and we predicted each cell’s corresponding gene expression profiles in the target species by appending the cell embedding with the target species factors. This approach allowed us to produce a denoised prediction of the gene expression profile of a given cell in the same or a different species. To make sure that the expression values are comparable across species, we normalized each cell’s gene expression profile based on 2,401 housekeeping genes in mouse [40]. Then we compared gene expression changes across species and, for each gene, we calculated the log2 fold change (log2FC) between the two species for each cell, and we took the median value across all cells as its overall log2 fold change. Genes were then grouped into three categories based on their evolutionary origin:

1. Genes located in the X-added region (XAR) genes were added to the X chromosome in eutherian mammals, i.e. these genes are X-linked in eutherian mammals (e.g., mouse) but are autosomal in metatherian mammals (on chromosomes 4/7) (opossum) and appear on chromosome 1 in chicken.
2. Genes located in the X-conserved region (XCR) are X-linked in both eutherian and metatherian mammals, but are autosomal (on chromosome 4) in chicken.
3. Genes located on autosomes in mammals and chicken.

To test whether genes in the XAR or XCR tend to become upregulated when becoming X-linked (i.e. lose one copy in males), we performed a one-sided, one-sample Wilcoxon signed-rank test with the null hypothesis that the median log2FC within that gene group is smaller than or equal to -1. To test whether XCU is stronger in mouse than opossum, we performed a one-sided, one-sample Wilcoxon signed-rank test with the null hypothesis that the median log2FC (between mouse and opossum) within that gene group is greater than or equal to 0. Similarly, to test whether the dosage of gene expression is fully compensated (i.e. each X-linked gene is upregulated by two-fold to achieve the same expression level that it had when it was autosomal), we performed a one-sided, one-sample Wilcoxon signed-rank test with the null hypothesis that the median log2FC (between mouse and chicken) within that gene group is greater than or equal to 0.

## 3 Results

### 3.1 Icebear accurately predicts cell type profiles across species

Icebear is a deep learning model that is designed to integrate cross-species, single-cell profiles (Figure 1A, Methods 2.5). Once trained, the model decomposes each observed cellular gene expression measurement into several components, corresponding to cell, batch and species. Prior to applying the model in a prospective fashion, we carried out several validation experiments to verify that the model works as intended.

First, we hypothesized that we could reduce or eliminate effects associated with species or batch factors by manipulating the model appropriately. To test this hypothesis, we trained the model on a public dataset derived from multiple species, and we assessed whether the model could eliminate the effect of species in the learned cell factors. Specifically, we used as input a dataset with primary motor cortex (M1) cells from both human and mouse [35], and we mapped genes across species via orthology [41]. The original dataset shows clear separation by species in the context of 2D visualization via UMAP (Figure 2A). We then investigated whether Icebear could be used to remove this species-specific effect. Accordingly, we plotted a UMAP visualization of the learned cell embeddings (Figure 2B). This visualization shows minimal segregation of cells by species, suggesting that Icebear can correct species-specific effects and align single cells across species. To further quantify the performance of cross-species cell type matching, we compared Icebear with SATURN, a state-of-the-art cross-species alignment method [14]. Although cell type matching is only a by-product of Icebear, we find that Icebear can outperform or perform comparably to SATURN in this task (Supplementary Figure S1). To further quantify the performance of cross-species cell type profile prediction, we created another baseline that is based on cross-species alignment [13] followed by *k*-nearest-neighbor calculation, and we found that Icebear significantly outperforms this baseline (Supplementary Figure S2).

**Figure 2:**
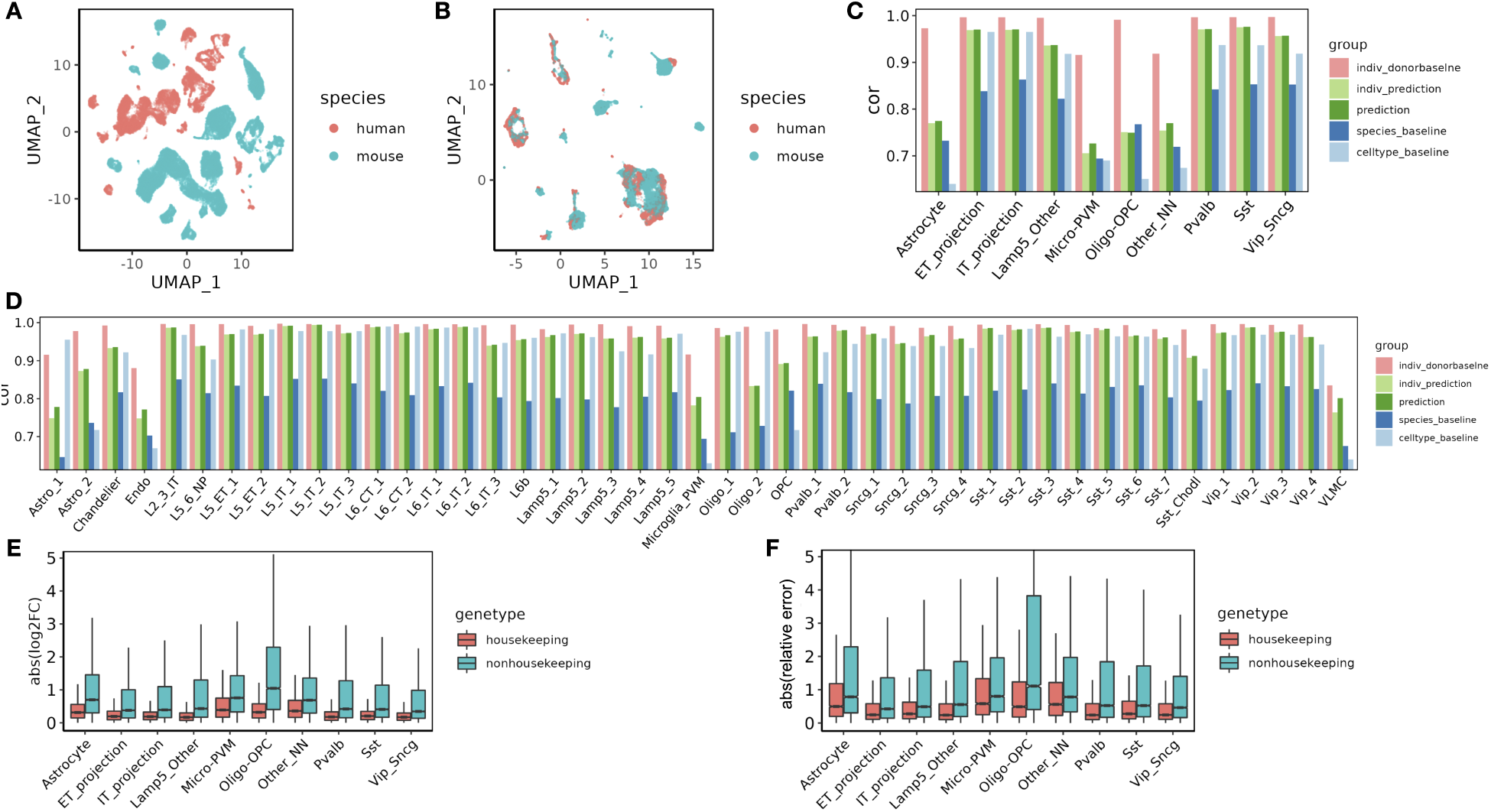
Alignment and prediction of single-cell profiles from mouse to human in primary motor cortex. (A) UMAP of cells across species, colored by species. (B) UMAP of cell embeddings learned in Icebear, where cell embeddings are independent of species factor. (C) Barplot showing Pearson correlation coefficient (cor) between predicted and observed gene expression profiles in the ”held-out” major cell types in human motor cortex. For each cell type, the Pearson correlation coefficient is compared between Icebear’s prediction (dark green bar) and two baselines (species and celltype). We also compared “indiv donorbaseline” (correlation between individual donors) with Icebear’s correlation with individual donors (“indiv prediction”, light green bar). (D) Similar to C, we evaluated Icebear’s performance on predicting high-resolution cell type specific profiles. (E) Boxplots of genes absolute log2FC between predicted and true human profiles per cell type (grouped by whether they are housekeeping genes or not). (F) Boxplots of genes relative prediction error across cell types.

The primary goal of Icebear is to jointly model gene expression data from human and mouse using factors representing species, gene, and cell identity, thereby enabling cross-species prediction and comparison. Ideally, the model could then be used to answer questions such as, “What would the expression profile of this mouse cell look like if it were instead a human cell?” Unfortunately, validating the accuracy of such a predictor is impossible. We therefore adopted an alternative validation approach.

To validate Icebear’s cross-species prediction performance, we made cross-species predictions at the level of individual cells but evaluated the predictive accuracy at the level of cell types. For this analysis, we used the cell type annotations produced in the original M1 study [35]. To mimic the real life scenario where we have uncharacterized biological contexts, we trained Icebear using a dataset in which one cell type in human was held out entirely, and we then used the trained model to predict gene expression profiles from the same cell type in mouse. Aggregating these predicted single-cell expression profiles yields a predicted pseudobulk profile that can then be compared, via Pearson correlation, to the pseudobulk profile of the held-out cells.

Before carrying out this experiment, we designed three “baseline” predictors to provide comparators for our model (Methods, Section 2.6). The first baseline (“donor baseline”) provides a “cheating baseline” on the predictive accuracy: we directly compute as our performance measure the mean Pearson correlation between pseudobulk profiles for the test cell type across all donors in the held-out test set. This baseline is cheating, in the sense that it has access to the data in the test set; however, the idea is that the empirical variance of the performance of this baseline on data from different individuals within a species provides a rough upper bound on how well any predictor could possibly perform on this task. The second baseline (“species baseline”) predicts that gene expression values do not change at all between species; thus, for a given cell type, the predicted human gene expression profile is equal to the corresponding mouse profile, with genes transferred from mouse to human through ortholog mapping. The third baseline (“cell type baseline”) is stronger than the species baseline, because it requires that we have access to cell type annotations in both species. In that setting, the cell type baseline makes a prediction for the held-out cell type by (1) finding the cell type in the source species whose pseudobulk profile most closely resembles that of the held-out cell type, and (2) identifying the corresponding cell type in the target species. The prediction for the held-out cell type is then the pseudobulk profile drawn from its neighboring cell type. Note that this baseline is somewhat unfair to Icebear, which is not given access to cell type labels in either species.

Validating on the cell types annotated in the M1 dataset, we find that Icebear outperforms both the species and cell-type baselines. We first applied the validation protocol to the 10 general motor cortex cell types, observing that in 9 out of 10 cases, Icebear outperforms both baselines (Figure 2C). The one exception is Oligo-OPC (i.e. Oligodendrocytes and Oligodendrocyte progenitor cells) in which our model only outperforms the cell-type but not species pipeline. To further evaluate how well our model can predict single-cell profiles with fine cell type resolution, we trained Icebear on the same dataset but held out and evaluated the prediction based on high-resolution cell type annotations (Figure 2D). In this setting, Icebear outperforms the species baseline in all 45 cell types and outperforms the cell-type baseline in 33 of the 45 cell types (*p* = 5.62*×*10*^−^*^5^, Wilcoxon one-sided signed rank tests). Note that our model tends to perform better in neurons than non-neuronal cells, especially endothelial cells, microglia-PVM (perivascular macrophage) and VLMC (vascular leptomeningeal cells). As expected, our method performs worse than the “donor baseline.” This suggests that though the model can perform general cross-species imputation, it may fail to capture some cell type-specific evolutionary effects.

Because Icebear performs non-linear projection of cellular profiles across species, we hypothesize that a gene with large cell-type specific functional divergence during evolution may participate in species- and cell-type-specific adaptation processes, and thus may not follow a general cross-species expression pattern shift. Thus, we expect that the expression values of functionally diverged genes may be poorly predicted across species. To test this hypothesis, we investigated whether housekeeping genes, whose functions are more likely to be conserved across species, can be more accurately predicted compared to non-housekeeping genes. To do that, we calculated absolute log2FC between the predicted and observed gene expression values in human (Figure 2E). Indeed, of all major cell types tested, Icebear achieved better predictive power for housekeeping genes compared to non-housekeeping genes. To further control for potential bias caused by baseline expression and expression differences across species, and to investigate how much Icebear’s prediction improves upon the species baseline, we calculated the absolute value of the ratio between the prediction error (predicted human vs. original human) and cross-species difference (original mouse vs. original human) (“relative prediction error”, Methods 2.6, Figure 2F). Again, Icebear is able to correct for species-specific effects more accurately for housekeeping genes than non-housekeeping genes. These results point to a potential limitation to any cross-species model, since the cell-type specific evolutionary divergences, which are a product of evolutionary selective pressure, may contribute to the gap between the predicted and observed profiles. To test whether the better predictive performance of housekeeping genes is due to higher gene expression, we divided all genes into 10 equal-sized bins based on pseudobulk gene expression across all cells and compared the predictive error of housekeeping versus nonhousekeeping genes within each expression bin (Supplementary Figure S6). The result suggests that genes with lower expression are likely to have a higher predictive error, and housekeeping genes tend to have lower predictive error than their expression-matched non-housekeeping genes. To gain insights into what factors drive such differences, we further hypothesized that genes with high expression variation across cells may be more challenging to predict than genes with relatively consistent expression. To test the hypothesis, we binned genes by expression variation across cells (Supplementary Figure S7). Intriguingly, while variance contributes to some performance differences, housekeeping genes still tend to have lower relative prediction errors than non-housekeeping genes. The above evidence suggests that Icebear’s prediction performance may reflect meaningful biological changes during evolution.

### 3.2 Icebear can generalize across tissues, datasets, and species

To test whether Icebear can generalize across tissues in another dataset, we trained the model on single-cell profiles from the human and mouse cell atlases [36, 37]. In this validation experiment, we focused on single-cell profiles collected in the adult stage (Methods 2.4.2). We held out each major cell type from human, and we trained Icebear to predict expression profiles based on the corresponding cell types in mouse. To ensure a fair comparison that considers tissue-specific variations, we evaluated the cell-type-specific profile prediction per tissue, even though such information is not used in our training.

The results of this experiment suggests that Icebear can outperform both the species-specific and cell-type-specific baseline in 34 cases out of 41 total (Figure 3A). Interestingly, Icebear outperforms the donor baseline in 22 out of 27 cases (when more than one donor exists for that tissue and cell type). Also, explicitly modeling tissue as a factor significantly improves the cross-species prediction performance (Supplementary Figure S3). These results suggest that Icebear provides a robust estimation of single-cell profiles across species and thus can be broadly applied to computationally impute single cell profiles in human based on measurements in mouse across tissues and cell types.

**Figure 3:**
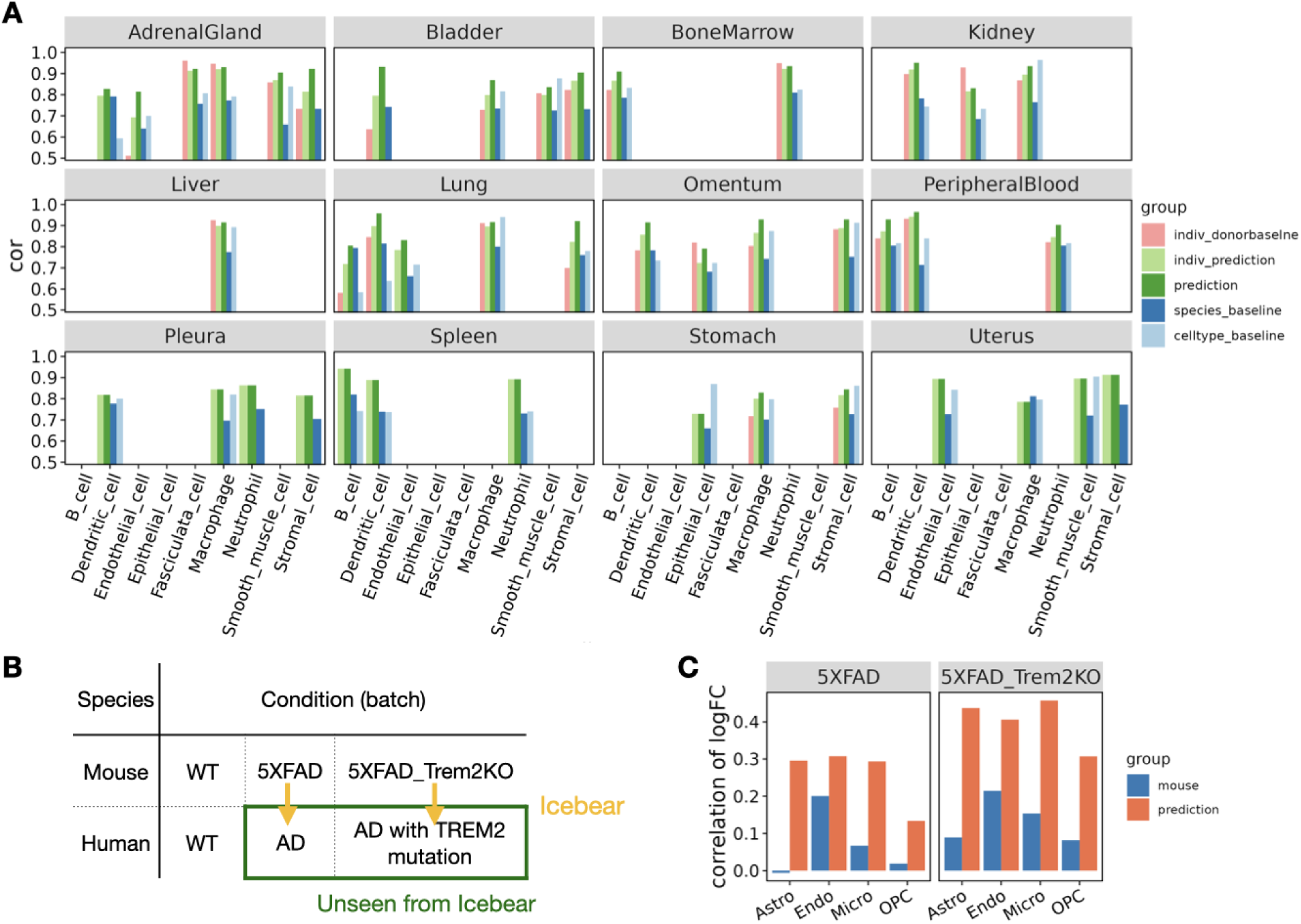
Prediction of human cell-type specific profiles across tissues and conditions. (A) Barplot showing Pearson correlation coefficient (cor) between predicted and true gene expression profiles in each held-out cell type per tissue. For each cell type, the Pearson correlation coefficient is compared between Icebear’s prediction (dark green bar) and the species and cell type baselines. We also compared “indiv donorbaseline” (correlation between individual donors) with Icebear’s correlation with individual donors (“indiv prediction”, light green bar). (B) Illustration of Icebear’s framework for predicting held-out human AD profiles from mouse AD models. (C) Comparison of predicted and true gene expression log2FC patterns in AD vs. WT. Barplots show Pearson correlation coefficients between observed and predicted log2FC in human (“prediction”, orange), and between original log2FC in human and mouse (“mouse”, blue).

We further applied Icebear to concatenated datasets of five species generated across four studies (Supplementary Methods S1.3). Icebear is able to align cells across the five species (Supplementary Figure S4A). Furthermore, Icebear outperforms all baselines when predicting human cell type-specific profiles from cells in the frog and zebrafish (Supplementary Figure S4B). These results further demonstrate Icebear’s ability to generalize to multiple species across various evolutionary distances.

### 3.3 Icebear can transfer findings from a mouse Alzheimer’s disease model to human

As a final validation and demonstration of a key use case, we test the hypothesis that Icebear can predict gene expression alterations in disease conditions versus healthy controls in human, based on healthy human samples and disease models in mouse. To test this hypothesis, we trained Icebear on human and mouse single-cell profiles in primary motor cortex (used in the above section), control samples in both human and mouse from an AD study, as well as profiles in the AD mouse model (Figure 3B) [38]. Once the model was trained, we applied it to predict the held-out profiles in human AD samples (green rectangle, Figure 3B) and calculated the log2FC of predicted gene expression between AD and WT samples in human. To evaluate how well the predicted gene alteration pattern agrees with the true pattern, we compared it against the true log2FC pattern derived from the original human AD versus control samples, using Pearson correlation (Figure 3C, orange, Methods 2.7). To make sure Icebear captures more informative disease signatures than the mouse model itself, we compared the true log2FC pattern in human against the log2FC pattern in mouse AD versus WT samples per cell type (Figure 3, blue). In all uniquely mapped cell types, Icebear’s prediction outperforms the mouse baseline, suggesting that by projecting single-cell profiles from mouse to human, Icebear is able to retrieve more accurate gene alteration patterns in human, compared to the original mouse experiments.

### 3.4 Icebear reveals X chromosome upregulation patterns during evolution

Having established the ability of Icebear to capture species-specific effects, we next apply the model prospectively, using it to investigate the pattern of expression change across species in specific classes of genes.

Essentially, our model allows us to ask, for any given cell, how its expression pattern would change if that cell had been in a different species. We are particularly interested in asking when autosomal genes become X-linked and have halved copy number in XY males during mammalian sex chromosome evolution, whether their gene expression increases from halved gene dose to compensate for this copy number changes. (Figure 1D, Figure 4C). To do that, we collected single cell RNA-seq profiles in male chicken, opossum and mouse brain samples, as well as male chicken and mouse heart samples (Figure 4A, Methods 2.4.2, with opossum heart not measured). We then applied Icebear on the dataset and predicted each cell’s gene expression changes when we swap its species factor across chicken, opossum and mouse (Figure 4B, Methods 2.8). Compared to the original measurement, Icebear is able to bring mouse and human cells closer to a shared space, as measured by the LISI score (Supplementary Figure S8). For each gene, we can do this analysis on a cell-by-cell basis, asking how its expression changes when the cell changes from, say, chicken to opossum or opossum to mouse.

**Figure 4:**
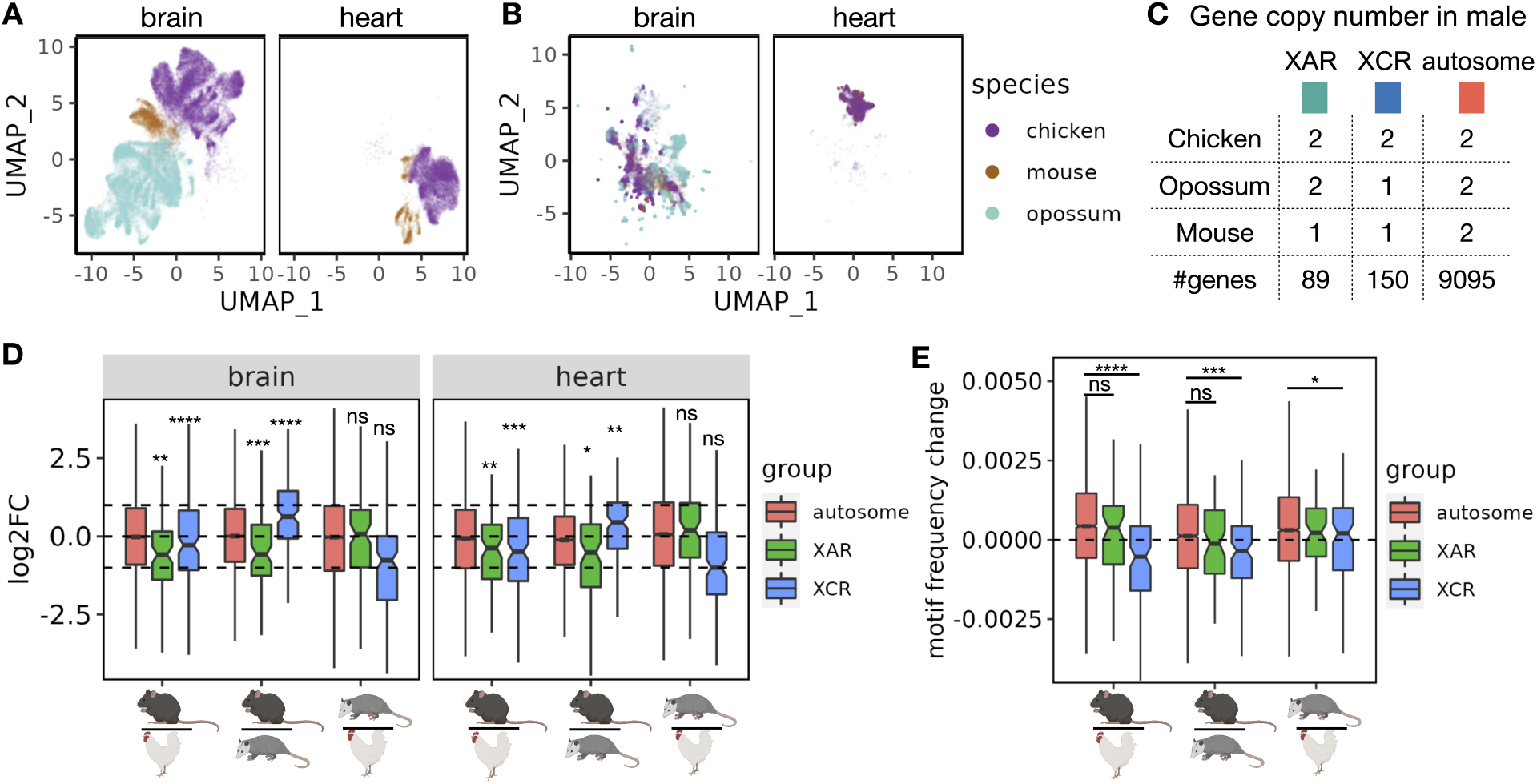
X chromosome gene upregulation pattern across species. (A) UMAP based on original input profiles. (B) UMAP based on cell embeddings learned from Icebear. (C) Gene copy number changes in males across species. (D) Boxplot of gene expression log2 fold changes from the lower to the upper species. Genes are grouped by their X-linked pattern. Statistical significance of XCU events are calculated using Wilcoxon one-sample rank sum test and subjected to multiple hypothesis correction. (E) Boxplot of GGACH motif frequency changes across species, grouped by the same rule as C. Statistical significance calculated on the hypothesis that genes in XAR and XCR are likely to have less m6A motif binding when moved from autosome to X chromosome, compared to genes in autosomal regions.

We find that when genes lying on the X conserved region (XCR) changed from the chicken autosome 4 (chr4) to the opossum X chromosome, the median gene expression log2FC between opossum and chicken do not fall significantly above or below -1 (*p* = 1.7 *×* 10*^−^*^1^ in brain and *p* = 1.3 *×* 10*^−^*^1^ in heart, Wilcoxon one-sample rank sum test, Figure 4D). Interestingly, these XCR genes show significant up-regulation on mouse X chromosome as shown by the median gene expression log2FC between mouse and chicken (*p* = 3.0 *×* 10*^−^*^7^ in brain and *p* = 4.8 *×* 10*^−^*^4^ in heart), suggesting that XCU of XCR genes is more prominent in eutherian mammals than in marsupials.

In addition, for genes at X added region (XAR) that changed from the chicken autosome 1 (chr1) to the X chromosome in eutherian mammals after the divergence from metatheria marsupials (e.g., opossum in which these genes on chr4 or chr7), we observed their median log2FC between mouse and chicken, and between mouse and opossum, are significantly more than -1, suggesting X upregulation also occurs for this group of genes in mouse (Figure 4D). Furthermore, there are no significant changes observed of XAR genes changing from chicken to opossum, when they are both on autosomes, which agrees with prior expectation (*p* = 6.1 *×* 10*^−^*^1^ in brain and *p* = 8.0 *×* 10*^−^*^2^ in heart). Comparing mouse with opossum or chicken, our finding also agrees with and further supports the general notion that X-upregulation does not usually fully compensate for the half-dosage effect [19], since XAR and XCR genes tend to have logFC less than 1 (in XCR regions when comparing mouse to opossum) or 0 (in other XCU comparisons) (adjusted *p ≤* 0.5 *×* 10*^−^*^2^ for all cases). Notably, Icebear is able to predict XCU patterns in heart, where single-cell measurements in opossum are unavailable.

A recent study has indicated that transcripts from X-linked genes including XAR and XCR genes tend to have lower levels of m6A (and thus less GGACH motifs) than those from autosomal genes, which results in more stable X transcripts compared to autosomal transcripts and thus contribute to XCU [26]. With this evidence, we hypothesize that the changes in m6A levels when genes move across species are related with whether the gene is in the XAR/XCR or autosomal. To test this hypothesis, and provide an orthogonal validation to XCU event, we retrieved the GGACH motif frequency in coding sequence (CDS) regions from chicken, opossum, and mouse [26], and ask whether there is a decrease in motif frequency for XAR/XCR orthologs when moved from chicken autosomes to mouse X chromosomes, compared to those from autosomal orthologs (Figure 4E). Indeed, we found that XCR orthologs show a significant decrease in the motif frequency between mouse and opossum or between mouse and chicken, when compared to autosomal orthologs. This agrees with Icebear’s prediction where XCU in the XCR is most significant in mouse (Figure 4D). In support of this, we only found marginal significance of motif frequency differences for XCR orthologs between opossum and chicken, in comparison to autosomal orthologs (*p* = 4.1 *×* 10*^−^*^2^). More intriguingly, we didn’t see significant changes in motif frequency for XAR orthologs compared to autosomal orthologs between mouse and chicken or between mouse and opossum, suggesting that XCU in the XAR may not be adapted through enhanced RNA stability via reduced m6A motifs in the CDS region.

## 4 Discussion

In this study, we proposed Icebear, a machine learning model for cross-species prediction and comparison of single-cell gene expression data. We demonstrate Icebear’s utility in predicting missing cell type-specific profiles between species, accurately transferring gene alterations identified in a mouse disease model to the corresponding human disease context, and identifying gene expression alterations during evolution in response to X chromosome dosage changes.

In our analysis, we have tested Icebear on both an atlas-level dataset covering 12 organs and 9 major cell types [36, 37], and a domain-specific dataset on motor cortex covering 10 major cell types and 45 fine-grained cell types [35]. Our results suggest that Icebear can perform accurate single-cell cross-species inference across various tissues and cell types. Interestingly, we observed that Icebear’s prediction accuracy varies by cell type. For example, the accuracy in predicting expression values of human non-neuronal cells on the M1 dataset tends to be lower than that of other cell types. There could be two reasons for this variability. First, previous studies have found that glia cells are more evolutionarily diverged [42], so a general cross-species algorithm trained on cell types other than astrocytes may not accurately capture astrocyte-specific variations. Cells with immune functions, such as microglia, may be especially variable among species due to different exposure to pathogens. The other possible reason is that neuronal cell types are more abundant and their profiles and functions are more cohesive than those of glial cell types, which could lead to an easier machine learning task. In line with this explanation, we observed that housekeeping genes are more accurately predicted across species, presumably because housekeeping genes tend to be functionally conserved across species and cell types and thus are more easily captured by Icebear’s projection framework. In the future, a closer look at the genes whose predicted expression profiles are very different from their observed profiles could potentially reveal genes that adapt to the species-specific environment in a cell-type-specific manner.

Icebear also shows improved accuracy at recapitulating transcriptomic perturbation patterns (based on log fold changes) in human AD based on mouse disease models, compared to traditional disease studies that directly map gene perturbation patterns across species by orthologs. This finding suggests a limitation of disease knowledge transfer through ortholog mapping, and demonstrates the potential of applying Icebear to more accurately transfer single-cell level perturbations such as disease signatures and drug responses from model organisms to human. In the future, we plan to develop statistical methods to rigorously assess the significance of predicted gene expression perturbations and compare them with the perturbation pattern based on original gene expression profiles, by leveraging ideas proposed by Boyeau *et al.* [43]. In this study, we demonstrated as a proof of principle how we can predict human AD profiles based on mouse AD models. We envision this method being applied to different diseases, such as Rett syndrome, Parkinson, lupus, or progeria where human samples (e.g. brain) are difficult to obtain.

Icebear reveals that there is an increase of expression levels of X-linked genes compared to their autosomal orthologs in mouse, supporting the hypothesis that XCU occurs during sex-chromosome evolution (Ohno’s hypothesis) [44]. The level of upregulation varies among individual X-linked genes, suggesting a gene-by-gene adjustment. Multiple transcriptional and post-transcriptional types of regulation have been suggested to explain XCU in mammals [23, 27, 45–48]. The most recently proposed regulatory mechanism is reduced GGACH motifs at X-linked versus autosomal transcripts, which results in X-specific reduced m6A levels and thus enhanced RNA stability [26]. Surprisingly, we found that XCR but not XAR genes show a significant decrease in frequency of GGACH motifs compared to autosomal genes in mouse, suggesting differences in XCU mechanisms dependent on the evolutionary origin of the XCR and XAR. Interestingly, X inactivation, which has been proposed to counteract XCU in females (reviewed in Disteche *et al.* [18]), is also more complete for genes in the XCR versus the XAR [49].

Icebear has the following advantages when compared with pseudobulk-based comparison (Supplementary Figure **??**). First, Icbear can discover XCR/XAR patterns when gene expression measurements are missing in certain species and tissues; i.e., we are able to predict the X chromosome upregulation pattern for the heart when there are no measurements available in the opossum heart. Second, Icebear’s cross-species comparison does not require any known matching of cell type labels across species or assumption of an equal proportion of cell type compositions between species, which greatly broadens the use cases where there are no existing matched cell type annotations across species. Third, comparing our prediction with the original measurements, we found general agreement of X chromosome up-regulation (XCU) patterns inferred from Icebear and pseudobulk profiles. However, XCU patterns are more apparent by Icebear imputation (as shown by increased significance) than simple pseudobulk comparison, because the latter may suffer from insufficient normalization cross-species and complex cell type composition. In addition, Icebear can predict XCU events when genes in the XAR region moved from the chicken autosome to the mouse X crhomsome, whereas the pseudobulk-based measurements cannot, again suggesting the sensitivity of Icebear. Icebear’s prediction agrees with the previous knowledge that mouse X-linked genes are subject to XCU and also suggests for the first time that the extent of XCU varies between regions with different evolutionary origins. In evolutionary biology, it is still controversial whether there XCU occurs in mammals and, if so, what and how X-linked genes are upregulated during sex-chromosome evolution. The challenge is the lack of appropriate data and computational methods to directly compare expression levels of orthologous genes before and after changing from autosomal to X-linked. Understanding X-chromosome dosage regulation is important to understand the impact of large-scale gene dosage alterations such as chromosome aneuploidy, which are often linked to cancer and developmental disorders. One of the major contributions of Icebear is its ability to compare gene expression changes across species at cellular resolution, so that we can reveal how orthologous gene expression evolves across different cells and tissues. In this work, we demonstrated that up-regulation of X-linked genes occurs in an evolutionary and diverse pattern and is, at least partially, associated with enhancing RNA stability. These findings provide new evolutionary insights into the time and mechanisms of XCU. In addition, by looking at the differences between the actual gene expression and the predicted value across species, we have observed the cell types and genes that have the most evolutionary divergence, which have the potential to explain what cell types and genes are conserved across species or adapted to species-specific needs.

Currently, Icebear is designed to model cross-species gene expression using one-to-one orthology relationships. This approach discards a subset of duplicated genes that potentially harbor valuable information about cellular function and evolution. In this study, we observe that the advantage of Icebear’s cross-species alignment performance against SATURN shrinks as the median cell UMI decreases (Supplementary Figure S1). In the future, inspired by ideas from SATURN [14] and coupled variational autoencoders [50], we could extend Icebear to include all genes, rather than only one-to-one orthologs. In principle, such an extension to Icebear could enhance the cross-species translation performance without losing information from non-orthologous genes. Icebear assumes the cell factor is conditionally independent of other factors (such as species and batch), according to the biological knowledge that cell identities are not supposed to vary a lot across batches or species. This assumption allows us to swap factors such as species to directly assess how the same cell’s expression would change across species. Note that in some instances cell populations may be distinct across species (e.g., mice have tails and humans don’t), which will not fit into our model.

In the future, we envision Icebear to be a general tool to (1) augment the effort of measuring complete profiles of human cells, (2) predict gene transcriptional changes in under-characterized human contexts by leveraging mouse models, and (3) study the evolutionary changes of transcription regulation, as well as divergence of cell types and genes. Icebear can also be extended in a straightforward fashion to perform cross-species prediction and comparison of other data modalities (e.g. protein quantity, epigenetic marks), when there are shared feature spaces across species.

## Data and code Availability

The Apache-licensed Icebear source code is available at https://github.com/Noble-Lab/icebear. We are working with the 4D Nucleome Consortium to make the sci-RNA-seq data publicly available.

## Supplementary information

### S1 Supplementary methods

#### S1.1 Comparison with SATURN

One of the key strengths of Icebear is its ability to predict and assess how a gene’s orthologs change expression across species. In particular, by making use of one-to-one ortholog mapping, Icebear can directly swap species factors to measure gene expression changes for orthologous genes during evolution. In contrast, SATURN uses macrogenes to perform cross-species cell alignment but not expression changes, and by design is incapable of predicting gene expression changes across species [14].

To compare Icebear with SATURN, we therefore can only evaluate the two methods with respect to cross-species alignment. To do so, we trained SATURN and Icebear separately on the primary motor cortex (M1) dataset [35] and human and mouse cell atlases [36, 37], because these two datasets have matching cell-type annotations across species. For each method, we used the LISI score to measure, for each cell type, how well cells from different species are aligned. The SATURN paper adjusts a single hyperparameter, the number of macrogenes, so we experimented with values of 1000, 2000, 3000, and 4000. At 5000 macrogenes, SATURN exceeded the available GPU memory (8 GB).

#### S1.2 Comparison with Harmony+kNN

We repeat the same protocol and evaluation of cell type prediction in the motor cortex dataset, where we hold out a cell type in human one at a time, run Harmony [13] to align the rest of the cells across species, retrieve the *k*-nearest human cells of each mouse cell from the held-out cell type, and use the pseudobulk profiles of all the *k*-nearest human cells as the baseline prediction. We tuned twice as many hyperparameters (12, rather than 6) for this method, including the number of principal components (PCs) used for Harmony (*n*=25, 50, 100), and the number of neighboring cells used to make the prediction (*k*=5, 20, 80, 320).

To select the best hyperparameters, we first choose the number of PCs based on cross-species cell alignment (calculated by the LISI score). We then select the best *k* based on a “cheating” method, where we pretend we already know the held-out cell type’s nearest neighboring cell type correspondingly in human and mouse, and we select the best model that gives the best pseudobulk Pearson correlation prediction on the neighboring cell type. This method creates an unfair baseline for Icebear, as Icebear does not leverage any cell type information during the training or model selection stages.

#### S1.3 Multi-species evaluation

To address the question of how Icebear works on more than three species and evolutionarily distant species, we constructed a five-species dataset, consisting of our in-house generated heart dataset collected from mouse and chicken, along with other public heart datasets from human [51]), frog (*Xenopus laevis*) [52] and zebrafish (*Danio rerio*) [53]. No shared batches exist across these four datasets, making it infeasible to perform cross-species comparison.

To test cross-species alignment and prediction, we first performed one-to-one orthology mapping following the protocol described in the manuscript (Methods 2.3). The frog dataset separately measures genes from the L (long) and S (short) homologous chromosome sets. To approximate one-to-one orthology mapping, we took the sum of gene expression counts across the two sets of chromosomes for each gene. Overall, 8675 ortholog groups were mapped across all five species. Meanwhile, without the transitive mapping step, only 7697 genes were included. We then retained orthologs with measurements across all five species and only retained the biggest batch from the human, frog, and zebrafish datasets. Next, we filtered out genes expressed in fewer than 50 cells and cells with fewer than 100 genes expressed. The above process ends up with 3064 genes and 36,227 cells.

Finally, we trained the Icebear model on this five-species dataset. In the first task, we used the default hyperparameters by setting the dimension of the embedding layer to 25 and ignoring the discriminator step, and then training Icebear on all cells. Using the resulting model, we asked whether cells from different species are aligned to the same space. As a control, we also generated cell embeddings derived from UMAP on the original measurement. In the second task, we identified shared cell type labels from species where cell type annotation is available (i.e., human, frog, and zebrafish), held out each cell type in human, and trained an Icebear model on the rest of the cells. We report the Pearson correlation between the original held-out profile and Icebear-predicted human cell type-specific profile based on the corresponding cell type in the frog or zebrafish (Methods 2.6).

### S2 Supplementary figures

**Figure S1:**
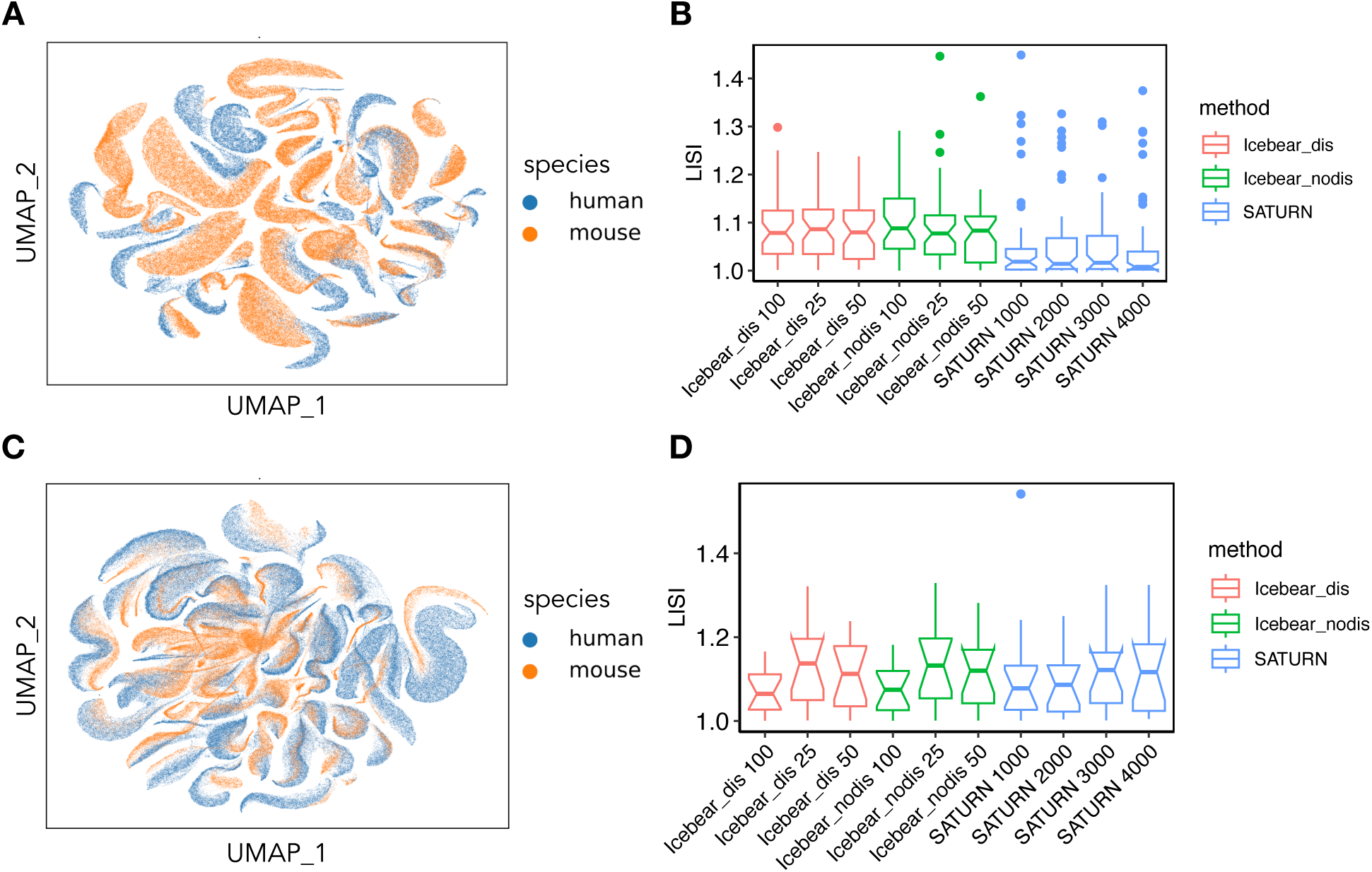
Icebear is comparable to or more accurate than SATURN at aligning single cells across species. (A) UMAP of SATURN-aligned cell embeddings of the motor cortex dataset, colored by the species each cell comes from. Median UMI across cells: mouse = 12060; human = 20056. (B) Boxplot distributions of LISI scores measuring how well mouse and human cells from the motor cortex dataset are aligned per cell type. All hyperparameters tuned are shown. (C) Similar to (A), UMAP of SATURN-aligned cell embeddings of the cell atlas dataset, colored by the species each cell comes from. Median UMI across cells: mouse = 688; human = 591. (D) Similar to (B), the LISI score distribution of cross-species alignment per cell type is measured on the cell atlas dataset.

**Figure S2:**
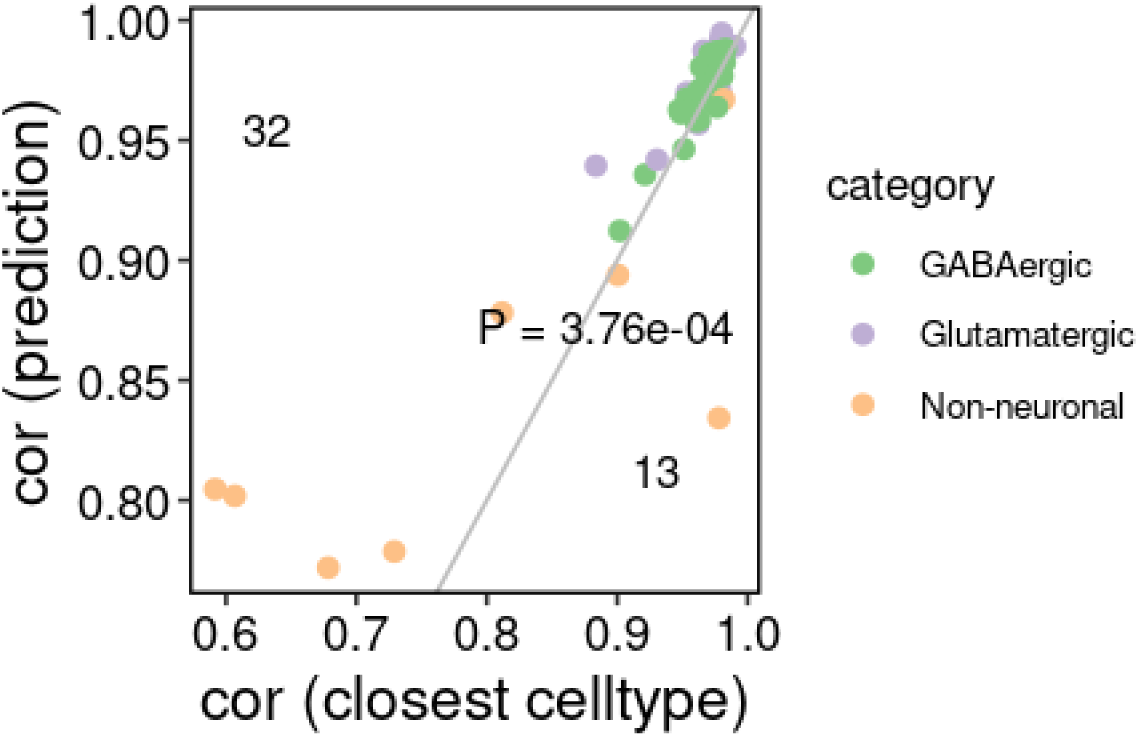
Comparison of Icebear and Harmony+kNN baseline. Pseudobulk Pearson correlation between the predicted and original profile in the mouse motor cortex dataset. Each dot indicates a cell type, colored by the major cell type annotation. The x-axis represents results generated by the Harmony+kNN baseline, and the y-axis shows results generated by Icebear. Numbers indicate the number of cell types off-diagonal. A p-value is calculated using Wilcoxon one-sided signed rank tests.

**Figure S3:**
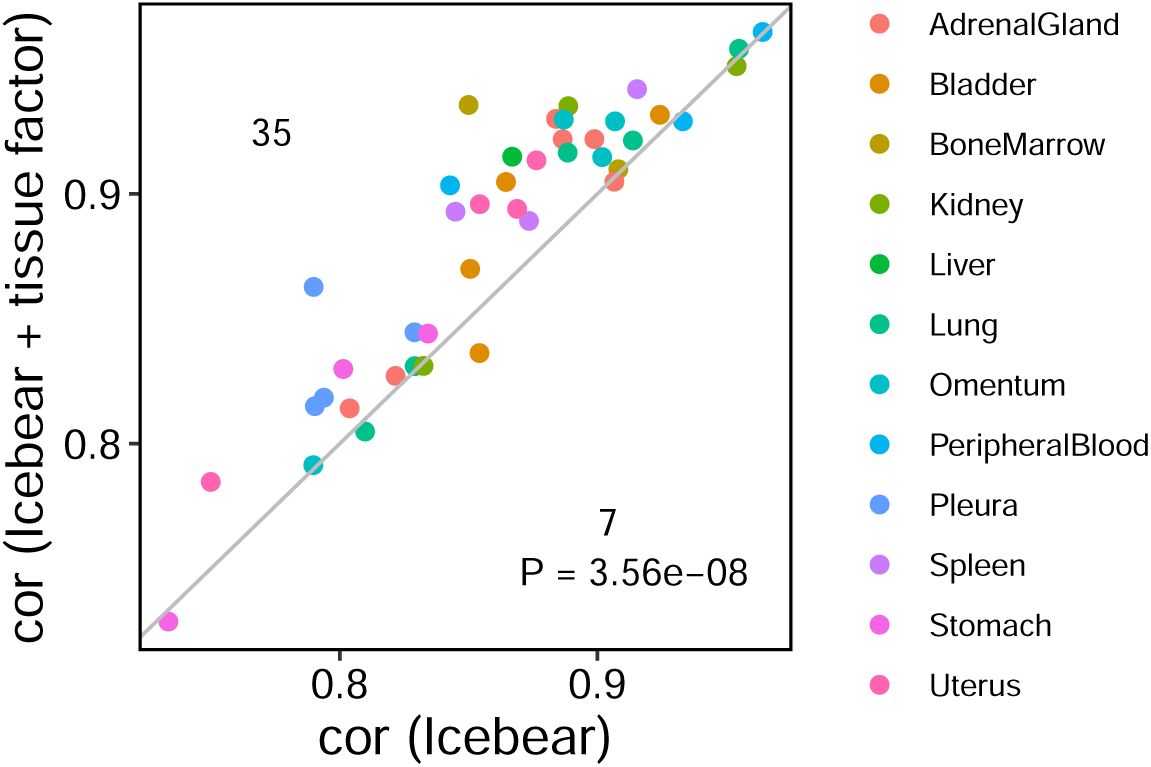
Adding a tissue factor improves Icebear’s performance. Pseudobulk Pearson correlation between the predicted and original profile in the cell atlas dataset. Each dot indicates a cell type, colored by the tissue it is extracted from. The x-axis represents results are generated by the Icebear model without the tissue factor, and the y-axis shows results are generated by Icebear with a tissue factor. Numbers indicate the number of cell types above and below the diagonal. A p-value is calculated using Wilcoxon one-sided signed rank tests under the null hypothesis that the median performance differences between tissue-aware and tissue-agnostic Icebear model is less than or equal to 0.

**Figure S4:**
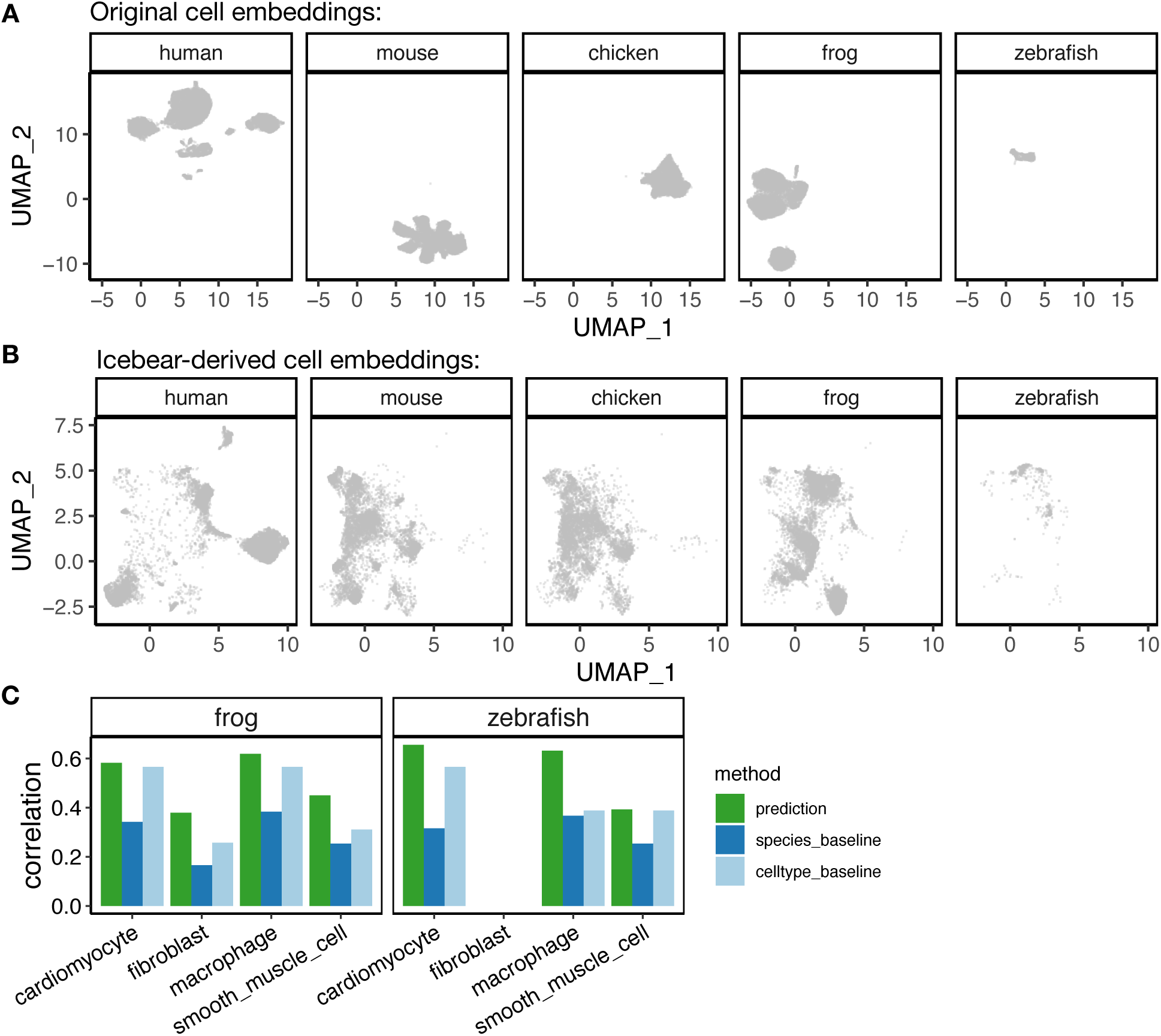
Icebear can be applied to multiple species. (A) UMAP of cell embeddings based on the original experimental measurement, cell embeddings are derived jointly and plotted separately across species. (B) UMAP of cell embeddings based on the Icebear model, cell embeddings are derived jointly and plotted separately across species. (C) Barplot showing Pearson correlation coefficient (cor) between cross-species predicted and observed gene expression profiles in the “held-out” major cell types in human heart. For each cell type, the Pearson correlation coefficient is compared between Icebear’s prediction (dark green bar) and two baselines (species and celltype). Human profiles are either predicted based on cells in the frog (left) or zebrafish (right).

**Figure S5:**
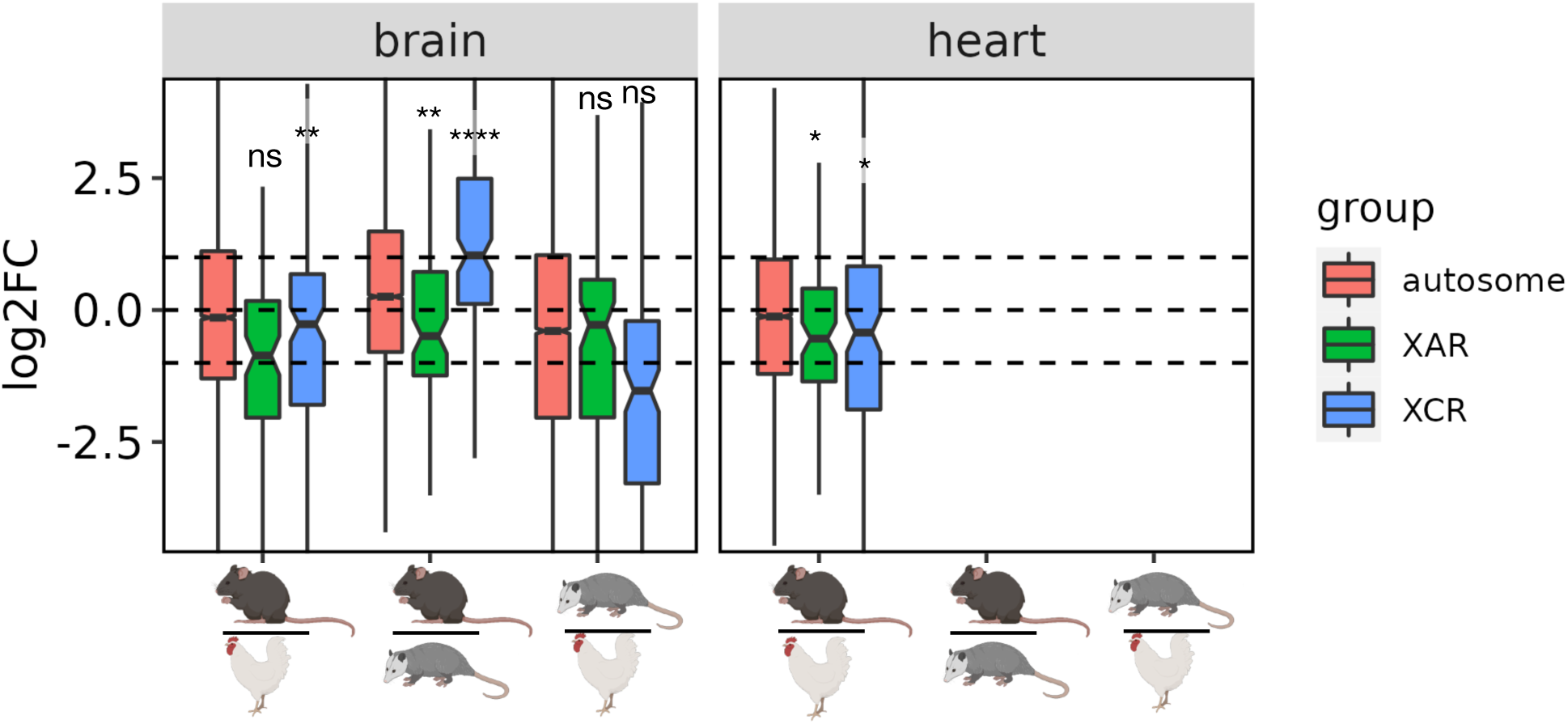
Comparison of XCU patterns between Icebear and pseudobulk measurements. Boxplot of log2 fold change of pseodobulk gene expression across species, with genes grouped by their X-linked pattern. The statistical significance of XCU events is calculated using the Wilcoxon one-sample rank sum test and subjected to multiple hypothesis correction.

**Figure S6:**
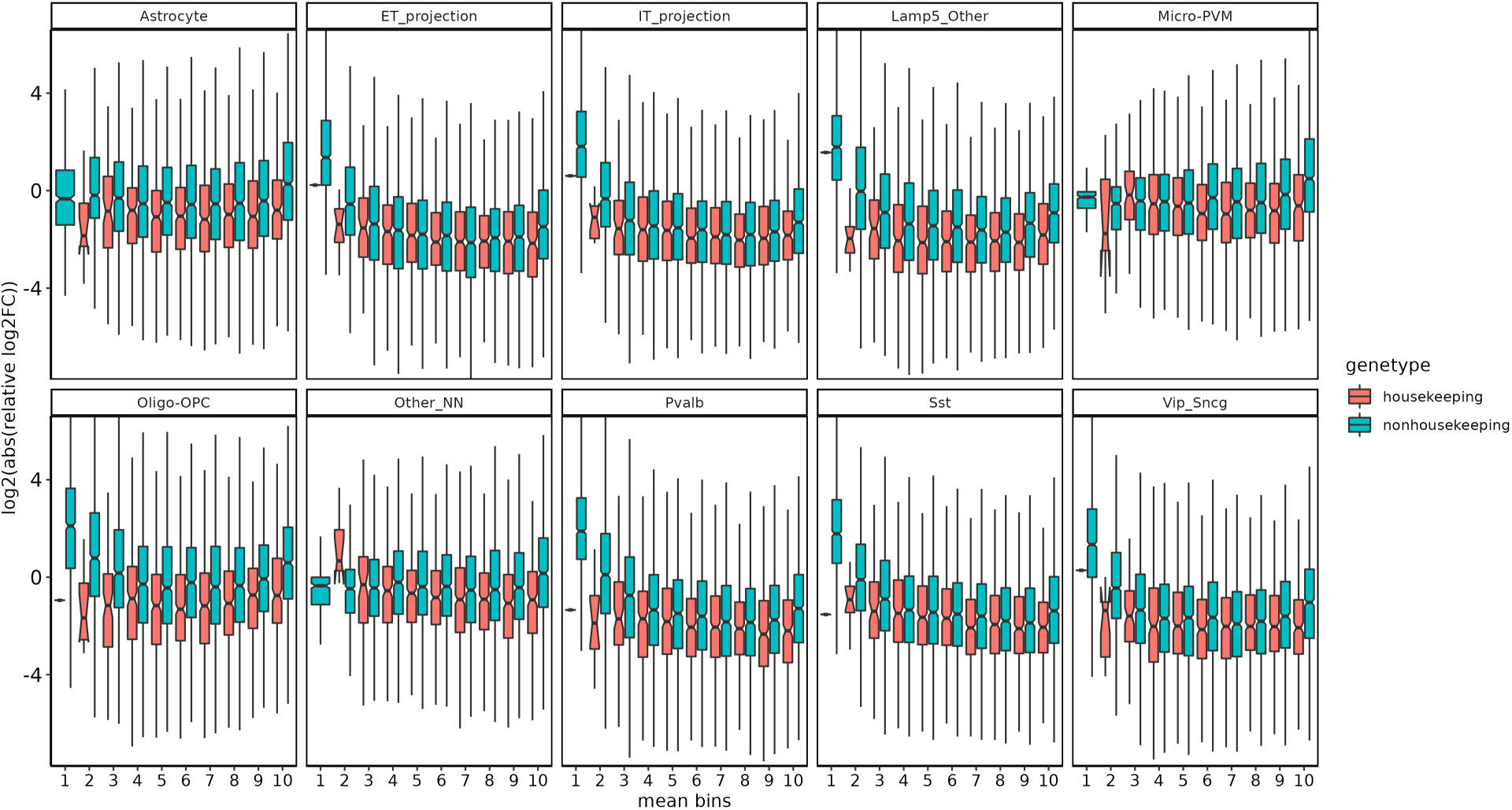
Relative prediction error across cell types and levels of pseudobulk gene expression. Boxplots of genes relative prediction error across cell types. Genes are separated by their properties, and ten equal-sized expression bins are sorted from lowest (bin 1) to highest (bin 10).

**Figure S7:**
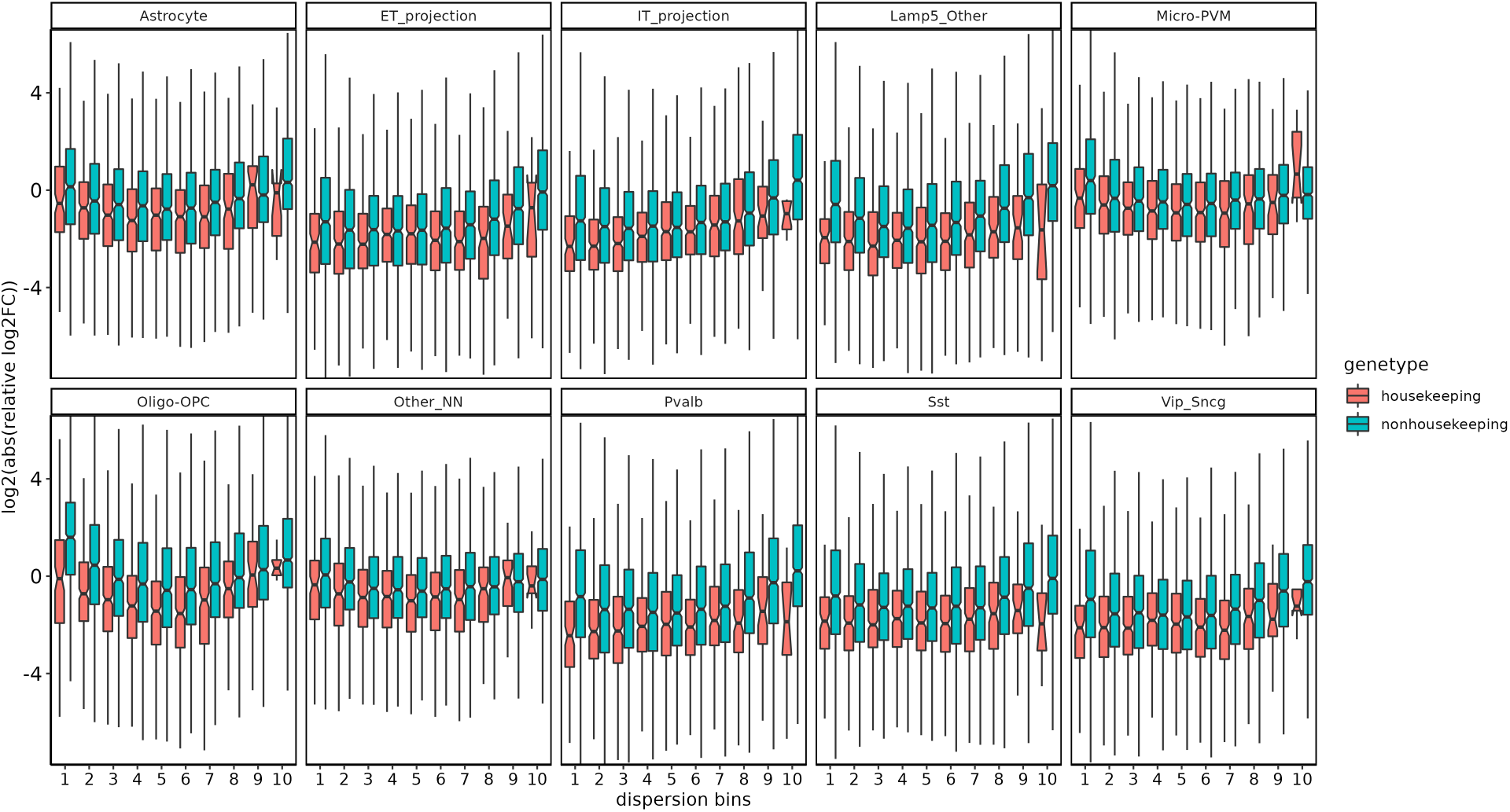
Relative prediction error across cell types and levels of gene expression dispersions. Boxplots of genes relative prediction error across cell types. Genes are separated by their properties and ten equal-sized bins based on their expression dispersion across all cells, from lowest (bin 1) to highest (bin 10).

**Figure S8:**
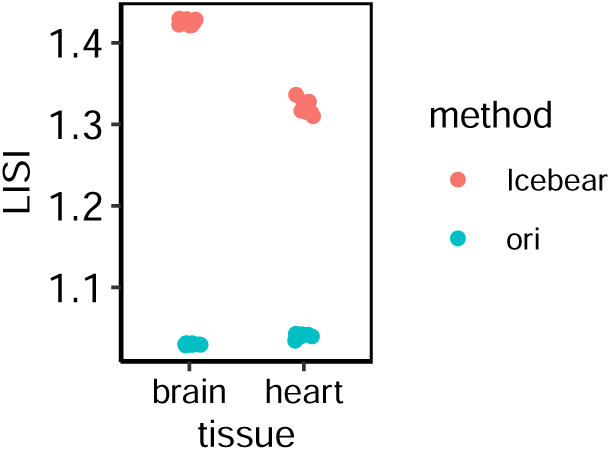
Quantification of cross-species integration on the sci-RNA-seq dataset. LISI scores calculated based on Icebear’s embeddings or the PCA embeddings of the original measurements. For each tissue (i.e., brain or heart), we calculate how well cells from different species are mixed together. 50,000 cells are randomly sampled from the cell population ten times, and each dot represents the LISI score on one random sampling.

